# Evolutionary patterns of 64 vertebrate genomes (species) revealed by phylogenomics analysis of protein-coding gene families

**DOI:** 10.1101/2020.03.31.017467

**Authors:** Jia Song, Xia Han, Kui Lin

## Abstract

**Background:** Recent studies have demonstrated that phylogenomics is an important basis for answering many fundamental evolutionary questions. With more high-quality whole genome sequences published, more efficient phylogenomics analysis workflows are required urgently.

**Results:** To this end and in order to capture putative differences among evolutionary histories of gene families and species, we developed a phylogenomics workflow for gene family classification, gene family tree inference, species tree inference and duplication/loss events dating. Our analysis framework is on the basis of two guiding ideas: 1) gene trees tend to be different from species trees but they influence each other in evolution; 2) different gene families have undergone different evolutionary mechanisms. It has been applied to the genomic data from 64 vertebrates and 5 out-group species. And the results showed high accuracy on species tree inference and few false-positives in duplication events dating.

**Conclusions:** Based on the inferred gene duplication and loss event, only 9∼16% gene families have duplication retention after a whole genome duplication (WGD) event. A large part of these families have ohnologs from two or three WGDs. Consistent with the previous study results, the gene function of these families are mainly involved in nervous system and signal transduction related biological processes. Specifically, we found that the gene families with ohnologs from the teleost-specific (TS) WGD are enriched in fat metabolism, this result implyng that the retention of such ohnologs might be associated with the environmental status of high concentration of oxygen during that period.

## 1. Background

With the recent advances in next-generation genome sequencing technologies, a large amount of high-quality genomes covering diverse taxa have been published[1]. The development and application of efficient and practical computational methods, such as comparative genomics[2], are very helpful for scientists to use these data to understand the underlying genetic mechanisms[3]. As one kind of comparative genomics strategies, phylogenomics[4] was firstly raised by Eisen JA in 1998. At first it had been exclusively defined as the prediction of protein functions from a phylogenetic view[4]. While in molecular systematics, phylogenomics is usually used to infer the evolutionary relationship of species using genome-scale sequencing data[5]. Uniting these two disparate definitions, phylogenomics is now widely regarded as the molecular phylogenetic analysis of genome-scale data sets[6], which can be used for predicting gene function[7-10], inferring evolutionary patterns of macromolecules[11-13], establishing the relationships and divergence times of genes/species[14, 15], exploring the genome duplications[16-19], and so on.

Phylogenomics data are available in several databases, such as EnsemblCompara[20], PhylomeDB [21] and Panther[22]. But high-quality phylogenomics data is still indispensable. On the one hand, these databases are known to contain many errors and uncertainties[23]. Directly using them in orthology detection or genome dynamics study could lead to erroneous results[24]. The causes of these errors are variable. As far as concerned, these databases considered little in the following two aspects: 1) the differences in histories of genes and species because of a hierarchy of evolutionary processes[25]; 2) the different selection stress on duplication/loss events in different gene families. On the other hand, most of these databases only contain data from model species, which is a limitation to the new sequenced genomes. Therefore, we believed an integrative and universal phylogenomics workflow, which is able to capture more differences among the evolutionary processes of different gene families and species, is imperative.

Here, we constructed a phylogenomics workflow mainly based on OrthoFinder[26], BEAST[27], Guenomu[28], RAxML[29], Notung[30], IQ-TREE[31] and SiClE[32] aiming to include the following two guiding concepts: 1) gene trees tend to be different from species trees but they influence each other in evolution; 2) different gene families have undergone different evolutionary mechanisms. In detail, an efficient species tree inference method and a parameter-learning method were proposed to model the evolutionary differences among different gene families and species trees. Based on protein sequences and CDSs (coding DNA sequences) from certain species, our workflow was designed to conduct species tree inference and duplication/loss dating following gene families’ classification and gene family tree inference/modification. As a case study, we applied our workflow to get the gene duplication history of 64 vertebrates’ genomes.

Duplications are of great significance as they would affect single gene, a stretch of several genes, whole chromosomes or even whole genomes and they are considered as the major driving forces for evolution of genetic novelty[33, 34]. However, many basic features of the evolution by gene duplication remain unknown[33, 35]. We applied our workflow on the genomic data of 64 vertebrates and 5 other eukaryotic species from Ensembl v84[36]. A species tree and 9,767 reconciled gene family trees were obtained. These results were then used to explore the WGD retention patterns and features, long-term local duplication preservation events and relative gene functions on vertebrate genomes.

## 2. Results and Discussion

### 2.1 An efficient gene tree–species tree phylogenomics workflow

#### 2.1.1 Introduction to our phylogenomics workflow

A phylogenomics workflow was constructed for multi-species genome evolutionary history exploration. As shown in Figure 1, the whole workflow could be divided into four processes. Under the guidance of the first guiding concept that we have mentioned above, the initial species tree was inferred based on the posteriors of gene families trees under a bayesian supertree model, which take both the gene duplication-loss and multispecies coalescent events into consideration. Meanwhile, inspired by supertree methods, whole genome-wide gene family trees were then used to revise the initial species tree based on the incongruent clades between the initial species tree and the available public species tree. In this way, it is able to efficiently reduce computational complexity by using the available species tree information and guarantee the accuracy by using genome-wide data. Then under the guidance of our second guiding concept, the fourth process in our workflow applied a parameter-learning process, which was designed to conduct gene tree modification and gene duplication/loss events dating. During the duplication/loss dating process, the parameters (event-costs: costdup and costloss) setting makes great influences[37]. In the previous studies[11, 20, 21], event-costs were usually set to the same values for all families. Here, we designed a parameter-learning process to find out the optimal parameter set for each gene family, which may help to capture the difference of selection pressures on gene duplication/loss in different gene families.

**Figure 1.**
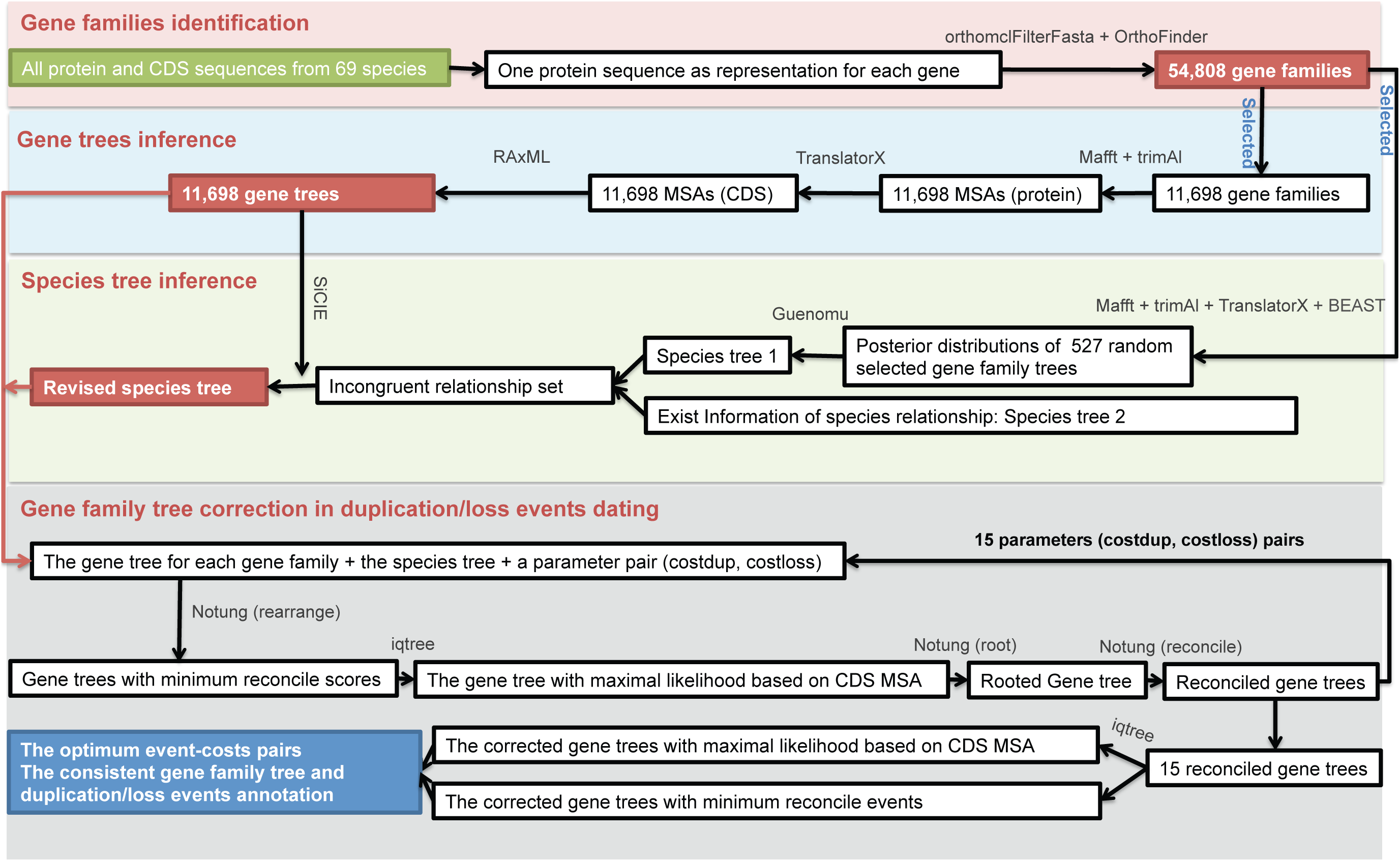
Flowchart illustrating our workflow. Our workflow mainly consists of four processes. The third and forth are the most important processes in our workflow. The inputs are displayed in green rectangles. The intermediate results are displayed in red rectangles while the final results are displayed in blue rectangles. The software, operation and some parameters used in this workflow are marked on the arrows in grey, blue and black font respectively.

### 2.1.2 Comparison with other similar works

In order to quantify the accuracy of our phylogenomics workflow, we compared the inferred species tree with the mammals species tree published by Song *et al.* 2012[38] (Figure S1 in additional file 1) and compared the inferred reconciled gene family trees with EnsemblCompara in ancestral genome content metric, ancestral chromosome linearity metric[24] and duplication consistency score[20]. Here, ancestral genome content metric is based on the assumption that the ancestral genome content sizes should be close to the extant genomes. Ancestral chromosome linearity metric assumed that each gene on ancestral genomes should have zero, one or two neighbors, with a peak at two while genes with three or more neighbors are the errors from the inferences. And duplication consistency score measures the intersection of the number of species post duplication over the union. It’s based on the assumption that most duplication should have the gene persisting at least in an equally likely manner in subsequent lineages[20].

Firstly, we compared the inferred final species tree (Figure 2) with Song’s mammals tree. Among the totally 31 shared species, only the tree shrew showed a incongruent evolutionary location between the two species trees. The correct location of tree shrew along the species tree is still under controversy[39]. Thus, our final species tree shows high accuracy in the mammals’ clade.

**Figure 2.**
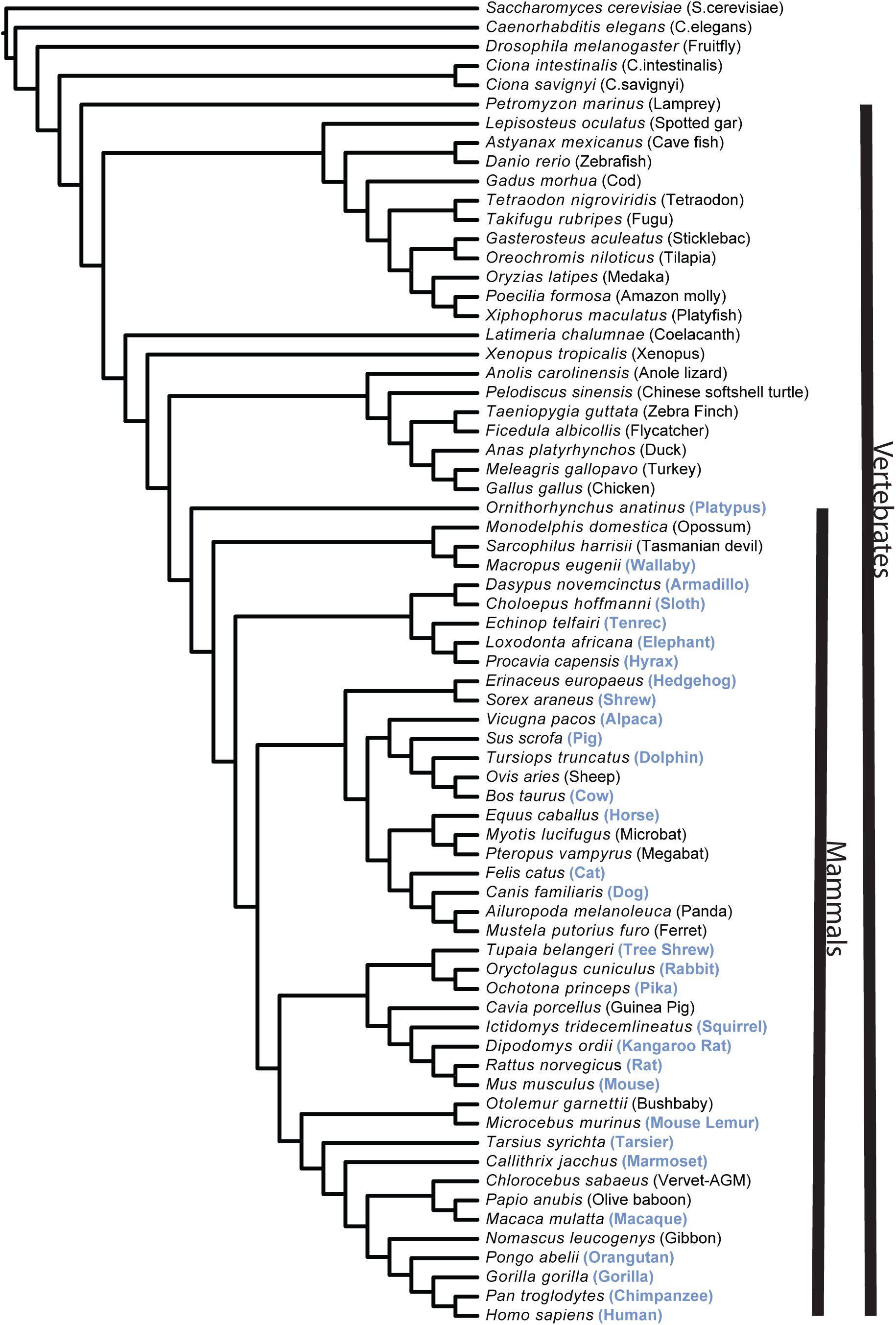
Final species tree. The common names of species are displayed in parentheses following the Latin names. And the common name of the common species between this species tree and the mammals species tree published by Song *et al.* 2012[38] are in blue font.

Secondly, according to our reconciled gene family trees, there were 50,916 duplication events occurred in the evolutionary history of 9,767 gene families. For the related 8,514 gene family trees from EnsemblCompara, there were 132,396 duplications. Then, as shown in Table 1, ancestral genome size inferred from our results shows closer average size to extant genomes than EnsemblCompara. As shown in Figure 3A, results from our workflow include much more ancestral genes with two neighbors and less genes with three or more neighbors compared with EnsemblCompara. Figure 3B shows clearly that the vast majority of the duplications from our workflow have a higher duplication consistency score compared with EnsemblCompara. Above all, EnsemblCompara output much more duplication nodes compared with our workflow. The vast majority of these duplications from EnsemblCompara perform worse on the three metrics mentioned above. Furthermore, we inferred another phylogenomics result by following our workflow but without the reconciliation parameter-learning in process 4. Results improved a little by the reconciliation parameter-learning according to the three metrics. Actually, the reconciliation parameter-learning process might have bring in more improvement on accuracy. Because we can only compare the results based on the 9,767 gene families in our core set while parameter-learning have already helped us to filter out the gene families which easy to receive wrong reconciliation results.

**Table 1.** Average genome sizes comparsion.

| Pipelines | Ancestral Genomes<sup>c</sup> | Extant Genomes<sup>d</sup> | Ancestral/extant Ratio<sup>e</sup> |
| --- | --- | --- | --- |
| Our pipeline<sup>a</sup> | 10290.25 | 9994.07 | 1.03 |
| Our pipeline<sup>b</sup> | 10457.22 | 9994.07 | 1.05 |
| EnsemblCompara | 22389.23529 | 13329.38 | 1.68 |

<sup>a</sup>Our standard workflow according to the processes as we described in Materials and Methods.
<sup>b</sup>Insteading of the optimal parameters with default parameters (1.5,1) when reconciliation by Notung in our workflow.
<sup>c</sup>Average ancestral genomes size according to our core data/Ensembl 8,514 gene families.
<sup>d</sup>Average extant genomes size according to our core data/Ensembl 8,514 gene families.
<sup>e</sup>Ratio of average ancestral genomes size and average extant genomes size.

**Figure 3.**
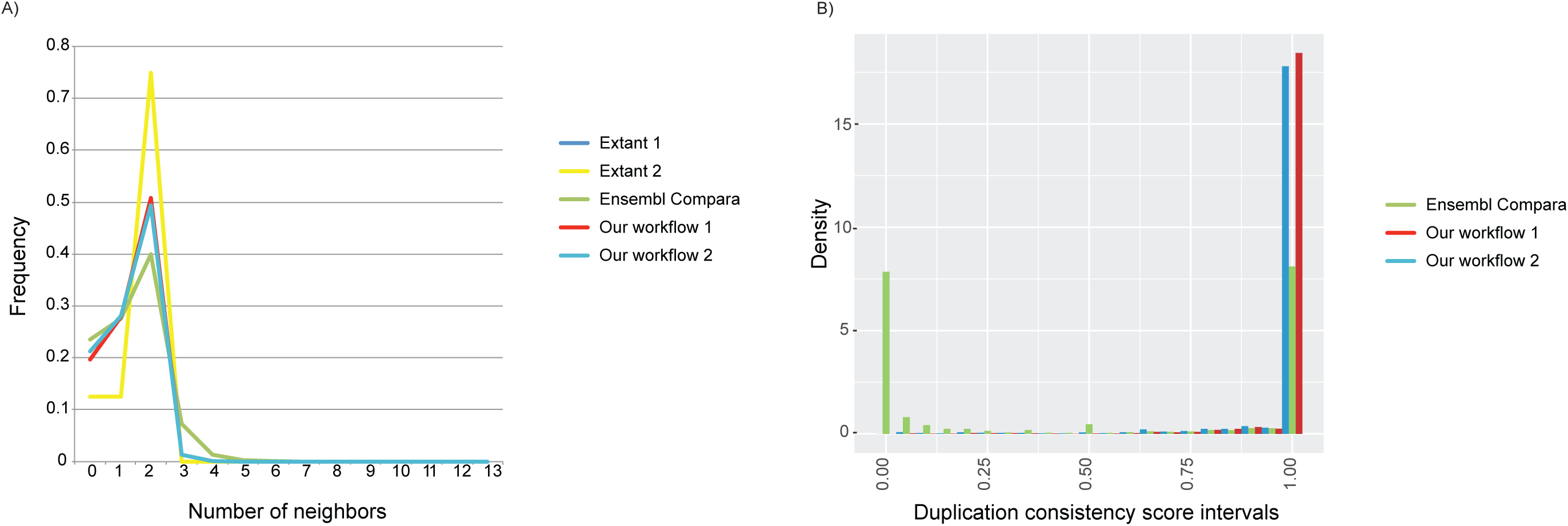
Comparison between our workflow and EnsemblCompara. ‘our workflow 1’ represents the standard workflow we have described in Materials and Methods section and ‘our workflow 2’ represents the same workflow but without parameter-learning in duplication/loss dating. **a**. Ancestral chromosome linearity metric. Extant 1 represents the genes neighborhoods status on extant genomes based on our 9,767 core gene families. Extant 2 represents the genes neighborhoods status on extant genomes based on Ensembl 8,514 gene families. The rest three represent the genes neighborhoods status on ancestral genomes inferred from different phylogenomics results. **b**. Duplication consistency score.

### 2.1.3 Limitations and future development

In the species tree inference process, only 527 gene families were used to infer the initial species tree to avoid costing too much computational time to get the gene family tree posteriors. Theoretically, most important information reflected by other gene families will be lost. So we revised the less supported clades on the initial species tree based on genome-wide gene family trees. However, there are two problems. First, algorithms that can deal with genome-wide gene families directly are more preferred. Second, there is no available species tree like the Ensembl species tree at most times.

Algorithms able to directly infer the species tree based on all gene families are more preferred. However, as the representations of the two main categories of such methods, Phyldog and *BEAST are not suitable for big scale family data. Firstly, methods as *BEAST cannot deal with paralogous genes which are common in gene families. Secondly, Phyldog[40] is limited by the sample size. Phyldog was designed to co-estimate genes and species trees under a DL model in a maximum likelihood framework, which get results in a short running time theoretically. Under our test, however, it was out of memory (our computational resource: 4T in memory) when we applied Phyldog on all the 11,698 families by default parameters. Then the family number was reduced to about 130, it can infer a species tree and 130 the gene family trees. From the Phyldog species tree (Figure S2 in additional file 1), we can see some obvious mistakes. Perhaps, the MSAs of the selected 130 gene families were not enough to reflect the real relationships of species. We will try to seek or develop an efficient species tree inference algorithm, which is able to co-estimate gene and species tree basing on genome-wide gene families for our workflow in the near future. Currently for our limited computational resources and large-scale data, our workflow may be a good choice. In addition, there is no available species tree like the Ensembl species tree at most times. To overcome this, it could be a proper way to choose two or more gene family sets randomly to get two or more initial species trees and compare the initial trees with each other to get the incongruent clades (Figure S3 in additional file 1).

In addition, read-through genes might also cause problems. A read-through/conjoined[41-43] gene is formed at the time of transcription by combining at least part of one exon from each of two or more distinct (parent) genes. In the gene family classification process, read-through genes/proteins will result in some nesting gene families including their parents. Such situations have not been considered in most phylogenomics datasets. In our case study, we seek these nesting gene families (Figure S4 in additional file 1) based on the read-through genes annotated in the GENCODE annotation file of HUMAN (V24) [44] and filtered out 454 such families. However, annotations of read-through/conjoined genes on other genomes are lacking or in low accuracy. It merits further attention to find a better way to deal with these families.

### 2.2 The features of whole genome duplication on vertebrate evolution

It is now clear that there have been three major WGDs in vertebrate genomes evolutionary history. Two (named 1R WGD and 2R WGD respectively) occurred near the base of the vertebrates’ evolutionary history and the third (named TS WGD) occurred at the base of the teleost fishes’ evolutionary history [45-49]. Although WGDs are often credited with great evolutionary importance, the processes governing the retention of ohnologs (paralogs generated by WGD) and their biological significance remain unclear. In this section, we explored the patterns of ohnologs retention and the relative function based on our reconciliation results of 9,767 gene families.

We got the gene families with ohnologs retention by seeking the duplications on the reconciled gene family trees and then mapped these duplications onto the species tree. Similar with previous studies[50-53], these three WGD-affected ancestral branches show about 9%∼16% gene duplication retention (additional file 2, supplementary material) which are significantly higher than other ancestral branches (P-value = 0.00193, Wilcox test, Table S1 and Figure S5 in additional file 1). Protein-protein interactions (PPIs) are enriched among the members of these gene families according to the human genes and their PPI data (Table S2 in additional file 1). This might reflect the gene dosage selection effects[54] after WGDs. Ohnologs retention from the 2R WGD might have undergone the weakest dosage selection among these three WGDs. Then based on duplication overlap rates (defined in Materials and Methods), we found that compared with other branches on the species tree, gene families with duplication retentions on the three WGD-affected branches are significantly more overlapped (P-value < 0.05,Fisher exact test). As shown in Figure 4A, 68 gene families retained ohnologs after all the three WGDs and 588 families after at least two WGDs.

**Figure 4.**
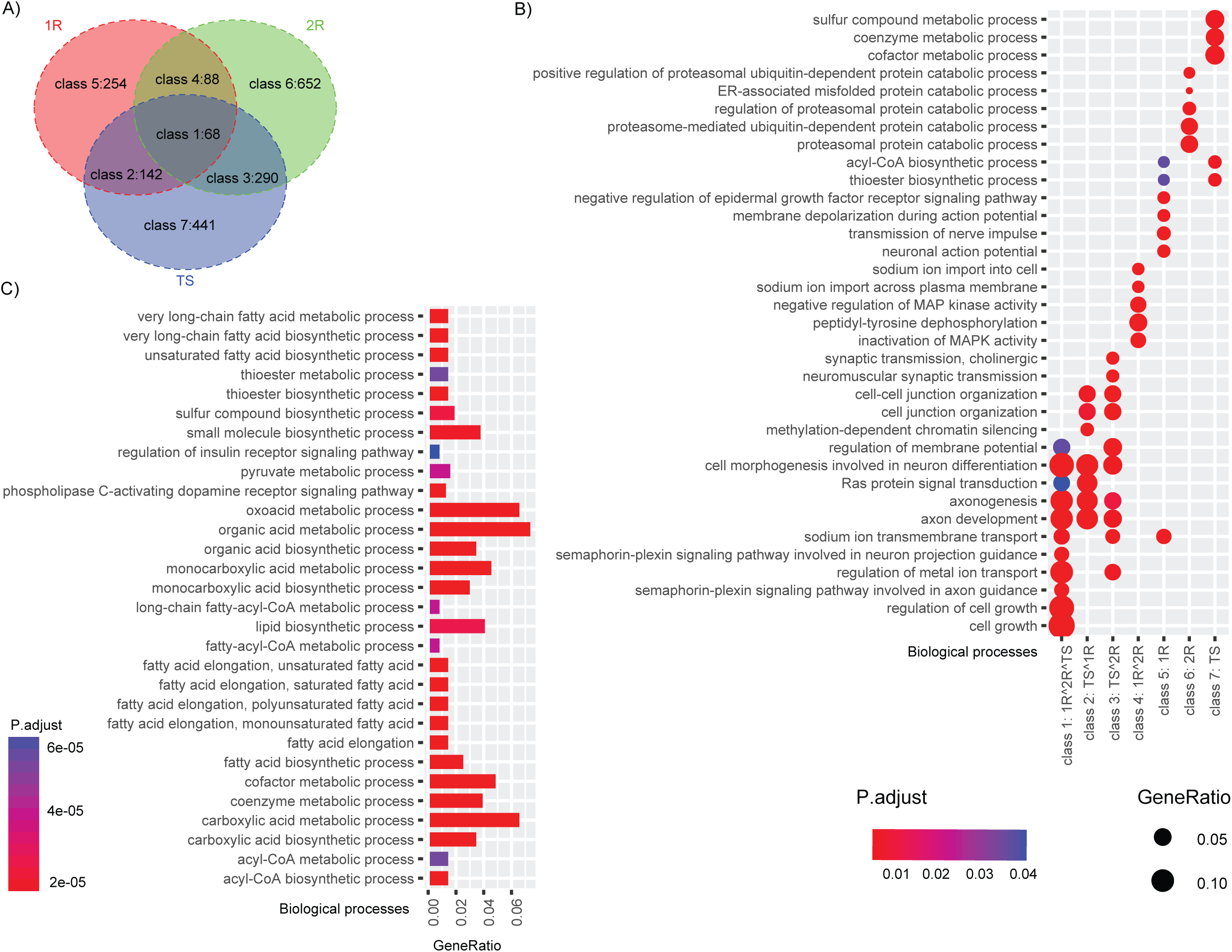
WGD-affected gene family classes and related gene function. **a**.Intersection among the three WGDs. Gene families with ohnologs retention are highly overlapped among the tree WGDs. We divided these ohnologs retention gene families into seven classes. 1R represents the first round WGD occurred on vertebrate genomes. 2R represents the second round WGD occurred on vertebrate genomes. TS represents the teleost fish specific WGD. **b**. Enriched functional categories comparison among the seven classes. The ‘A^B’ represents the intersection of ‘A’ and ‘B’. **c**. The biological processes enrichment results of class 7 which consist of gene families with ohnologs retention after TS WGD only. This analysis conducted based on gene ontology data of zebra fish.

According to human gene ontology information, gene families with ohnologs retention after these three WGDs are mainly involved in development, signaling and gene regulation (Figure S6 in additional file 1), which are consistent with the previous studies[55-59]. Then we divided these families into seven classes according to their ohnologs retention pattern after the three WGDs (Figure 4B). We found the 68 gene families with ohnologs retention after all of the three WGDs (class 1) are mainly involved in functional categories related to neuron, axon, signal and cell growth. Class 2 consists of gene families with ohnologs retention after both the TS and the 1R WGD and class 3 consists of gene families with ohnologs retention after both the TS and the 2R WGD. These two classes show similar GO enrichment results and they are both enriched in functional categories related to neuron, axon and cell-junction. Class 4, which consists of gene families with ohnologs retention after both the 1R and the 2R WGDs, are mainly involved in signal transduction. Above all, combined with the results from published studies[60, 61], nervous system and signal transduction related gene families are highly expanded on all vertebrate genomes through these three WGDs. Combined with the PPI enrichment results, this retention pattern may be a result of gene dosage selection.

The other three classes that consist of gene families with all ohnologs from one WGD are enriched in different functional categories and might reflect different retention mechanisms. We found that gene families in class 7 are enriched in fat metabolism. Further, we used the gene ontology data of zebra fish to redo the GO enrichment analysis, and the results (Figure 4C) showed more GO terms involved in fat metabolism, including anabolism and catabolism. As is well known, fat releases much more energy than other nutrients such as carbohydrate and protein in exhaustive oxidation and this process costs much more oxygen at the same time. More specifically, acyl-CoA and fatty-acyl-CoA, which included in many enriched biological processes, are essential products in metabolic process with oxygen consumption. Interestingly, this TS WGD happened at the period that the earth has its highest content of oxygen level (up to 33%) during the evolutionary history of vertebrates (Figure S7 in additional file 1). All of these lead to a suggestion that the high content of oxygen might be a kind of selection to the duplication retention after the TS WGD to promote fat as a main way to store energy. This might be one reason that fish have more unsaturated fatty and it is worth more discussion in future works.

### 2.3 The features of local duplications on vertebrate genomes

It should be noticed that many paralogs in current gene families were not originated by the WGD events mentioned above, but by extensive local duplications[62]. So we also identified the local duplications from our reconciliation results to explore such retention pattern in vertebrates. We firstly found that there were many more duplications occurred on the extant species-specific branches than the ancestral ones on the species tree (Kolmogorov-Smirnov test in R, p-value < 2.2e-16, Table S1 in additional file 1). As previous studies[63, 64] indicated that three steps are responsible for the generation of preserved gene duplications: origin through mutation (duplication), a fixation/spreading phase and a preservation/maintenance phase when the fixed change is maintained. The majority of duplications on extant species might still be under the fixation/spreading phase. While most duplications on ancestral genomes might already be under the preservation/maintenance phase for the most recent ancestral genome on our species tree existed 6.5 million years ago, which has already exceeded the average half-life of a gene duplication (approximately 4 million years) provided by previous studies[33].

Previous studies always focused on the duplication mutation rates and duplication fixation rates. Different from these studies, we estimated the duplication preservation/maintenance rates based on the duplications annotated on the ancestral genomes and their origin time (see Materials and Methods). After removing the duplications resulted from WGDs, we estimated the duplication preservation rates for 9,581 gene families. 7,075 gene families have no local duplications on ancestral genomes, which indicated that about 74% gene families in our core data have no long-term duplication preservation and most gene families kept singleton status during the evolutionary history of vertebrate genomes. We then got non-zero duplication preservation/maintenance rates for 2,506 gene families and 95% duplication preservation rates of these gene families are distributed between 0.0009 and 0.016 (Figure S8 in additional file 1). According to the gene and GO information from human, the gene families with long-term local duplications preservation are mainly involved in ion transport and some important signaling pathways. In addition, we also found that some local gene duplications might be retained through the natural selection caused by oxygen-level changes (Figure 5).

**Figure 5.**
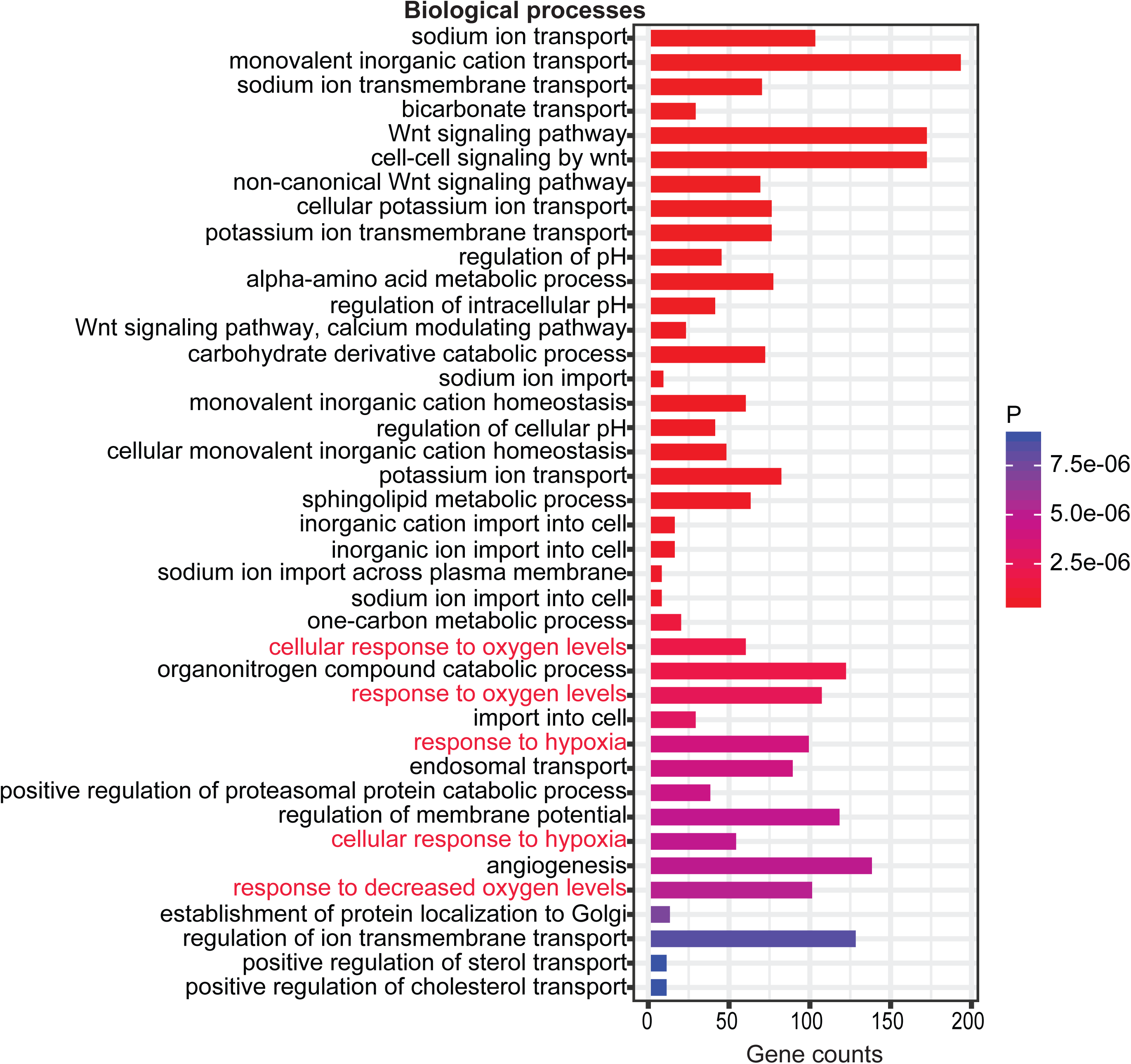
GO enrichment results of gene families with long-term local duplications retention. The oxygen levels response related biological processes are labeled in red font.

## 3. Conclusions

Based on two guiding concepts, we developed an integrative phylogenomics workflow by integrating an efficient species tree inference workflow, which adopt advantages from co-estimation and supertree methods, and a parameter-learning process to account for more about the relationship and differences among species and gene trees. It was designed for gene family classification, gene family tree and species tree inference and duplication/loss dating. Then, we analyzed the genomic data of 64 vertebrates and 5 out-groups from Ensembl as a case study to demonstrate a complete application of our workflow on the accurate inference of the evolutionary history of genome-wide gene families and species. Based on our phylogenomics results, we captured evolutionary traces from two different duplication retention mechanisms. We found that dosage selection might play an important role on ohnologs retention after WGDs and the changing environmental oxygen content might be a kind of natural selection affecting paralogs from both WGDs and local duplications. Above all, we expected that our workflow will facilitate further studies aiming to explore genome evolutionary histories.

## 4. Methods

### 4.1 Gene family classification

In order to get genomic sequences and annotations with high quality, we used the data of 69 species from Ensembl v84 as a case study to introduce our workflow. We downloaded all protein sequences and CDS sequences of these species from FTP site of Ensembl (http://www.Ensembl.org/, Build 84)[36] and chose their longest protein and CDS to be the representation for each gene. The too short (shorter than 10aa) and too simple (stop codons percent greater than 20) genes were filtered out then.

Here, OrthoFinder-0.4[26] was used to identify homology relationships between these sequences. OrthoFinder is a very efficient algorithm, which can overcome gene length bias and phylogenetic distance problems in gene family classification. After this step, we got totally 54,808 gene families. We then removed too simple (members from a unique species) and too complex gene families, which including known read-through genes (according to gene annotation file (v24) of human in ENCODE[44]) or with more than 1,000 members. 17,025 gene families were left for following analysis (additional file 2)

### 4.2 Gene family tree inference

In this step, protein sequences of gene families were aligned in MAFFT v7[65](--auto) and then translated into CDS alignments by translatorX[66]. The poorly aligned regions were removed from these CDS MSAs by trimAl[67]. Here, we removed some gene families with specific labels in its sequences (such as X) or with very poor alignment quantity (additional file 2). For the left 14,037 CDS MSAs, we inferred the gene family trees in RAxML v8.2.9[29] under GTRGAMMI sequence evolution model. For some MSAs including less than four members and some MSAs including too much gaps, we finally only got reliable phylogenetic trees for 11,698 gene families.

### 4.3 Species tree inference

There are 579 gene families including members from all of the 69 species. We filtered out the gene families with members’ distribution various largely (CV>0.5) on different species to avoid information asymmetry. So 527 gene families were left for species tree inference.

BEAST[27] (parameters: a gamma-distributed model of rate variation with four discrete categories and an HKY substitution model with a strict clock, 10,000,000 generations, sampling every 5000 generations) was used to infer the posterior distributions of these 527 gene family trees. The results possessed a good convergence under these parameters setting (with effective independent sample size greater than 200 for each parameter). Guenomu was used to infer species tree by considering gene duplication, loss and multispecies coalescent simultaneously (10,000,000 generations, sampling every 10,000 generations) based on these tree posteriors. It outputted two species trees and we used the one with 99.9% probability as the initial species tree.

In addition, we downloaded the Ensembl species tree inferred by EnsemblCompara to find out the possible errors on the initial tree. Firstly, we compared the initial species tree with Ensembl species tree to find out the incongruent clades. We found seven species (including ancestral species) bearing different phylogenetic sister-branches between these two trees (Figure S9 in additional file 1). Secondly, in order to find out the true phylogenetic sister-branches of these seven species, SiClE v1.2[32] was used to extract phylogenetic supports from the 11,698 gene family trees. The results (Table S3 in additional file 1) show that three clades on initial species tree got significantly higher supports than the respective clades on the Ensembl species tree. Conversely, other three clades on Ensembl species tree got significantly higher supports. Unfortunately, the rest incongruence couldn’t find a clear relationship from these 11,698 gene family trees. We then improved the initial species tree by modifying its three weaker supported incongruent clades. The final species tree is displayed in Figure 2.

### 4.4 Species/gene trees reconciliation

Inspired by the “Felsenstein equation” [68], we put forward a parameter-learning method to find out the optimal event-costs for each gene family based on two optimal principles. Firstly, the modified gene family tree should have largest ML (maximum likelihood) value based on the corresponding MSA (multiple sequence alignment) of CDS. Secondly, the optimal reconciled results should contain the fewest number of events to explain the incongruences between the gene family tree and species tree. Then, based on the optimum event-costs pairs of each gene family, we modified the low supported clades on the gene family tree and further dated the evolutionary events (duplication and loss) by reconciliation. In our case study, 11,698 gene family trees were used as inputs. Finally, 9,767 gene families got uniquely reconciled gene family tree under their optimal event-costs pairs. More details are described below.

We used Notung v2.8.1.7, IQ-TREE v1.5.2 and our parameter-learning scripts to finish species/gene trees parameter-learning and reconciliation. After the event-costs pairs (costdup and costloss) assignment, Notung is able to modify the gene family tree and date gene duplications/losses under a DL model in a parsimony strategy. In order to seek the optimal event-costs pairs set for each gene family, 15 event-costs pairs (costdup, costloss) with different cost ratios (costdup/costloss) were used to parameter-learning and reconciliation in a cycle process (Figure 1). The detailed steps are described as follow:

#### Step 1

Rearrange the gene family tree: the gene family tree was rearranged under the ‘Rearrange mode’ of Notung. We rearranged the weakly supported regions (edges with bootstrap less than 50) in the gene family tree to produce alternate gene family trees with minimum DL score based on the current event-costs pair. Here, at most 100 eligible alternate gene trees will be outputted. IQ-TREE was then used to pick out the most optimal one with maximal maximum likelihood based on the respective CDS MSA.

#### Step 2

Root the gene family tree: the gene family tree was rooted under the ‘Rooting mode’ of Notung by minimizing DL score based on current event-costs pair.

#### Step 3

Reconcile the species/gene tree: duplication/loss events were assigned on the gene family tree under the ‘Reconcile mode’ of Notung by minimizing DL scores based on current event-costs pair. Then current event-costs pair was set to the next pair and the analysis jumped to step 1 if the current event-costs pair wasn’t the last one. Otherwise, analysis jumped to step 4.

#### Step 4

Construct the optimal event-costs pairs set for each gene family: we used IQ-TREE to calculate the ML (maximum likelihood) for each resulted gene family tree, which was inferred under different event-costs pairs. In this way, we obtained the optimal event-costs pairs set I, which consists of event-costs pairs resulting in maximal ML trees. Meanwhile, we constructed the optimal event-costs pairs set II, which consists of event-costs pairs resulting in minimal DL events in the reconciled results. The intersection of optimal event-costs pairs set I and II was considered as optimal event-costs pairs III. The final optimal set for the family was empty if the optimal event-costs pairs set III contains more than one member and the reconciled results are inconsistent under members. Otherwise, the final optimal event-costs pairs set is the optimal event-costs pairs set III.

In our analysis, we totally used 15 (costdup, costloss) pairs (additional file 2). Finally, we obtained optimal reconciliation results for 9,767 gene families, which called core gene families in this study.

### 4.5 Comparison to EnsemblCompara

We downloaded the gene family trees inferred by EnsemblCompara from its FTP site. For our workflow integrated gene family classification, our gene families are not consistent with Ensembl’s. Here we selected 8,514 Ensembl gene families with more than four gene members and overlapped with the 9,767 gene families in our core results to do the comparison.

Based on the gene adjacencies extracted from annotation files of the 69 extant species, DeCo[69] was used to infer the ancestral genome contents and ancestral gene adjacencies according to our gene trees and Ensembl gene trees, respectively. Then, we calculated the duplication consistency score[20] for each duplication on these two gene tree sets.

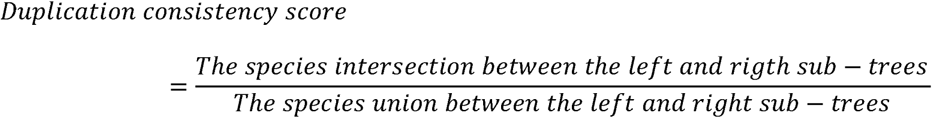

### 4.6 Others

#### 4.6.1 PPI and GO enrichment analysis

We used protein-linked information from STRING (v10.5) to finish the PPI enrichment analysis. We firstly abstracted the human protein-protein interaction network with combined score greater than 700. Then we abstracted the sub-network, whose nodes consisting of genes in our core gene families and edges linking members from different gene families. We found that there were 118,028 edges out of 68,641,957 gene pairs from different gene families on this sub-network. Then, we counted edges and such gene pairs among different WGD-affected gene family classes. As Table S2 (additional file 1), we found the PPIs were enriched in these classes (Fisher exact test).

The GO (gene ontology) enrichment analysis in this study was conducted by R package named ‘clusterProfiler’[70] basing on annotation data from ‘org.Hs.eg.db’ and ‘org.Dr.eg.db’.

### 4.6.2 Local duplication preservation rate

The local duplication preservation rates were inferred based on the gene duplications on ancestral genomes and their origin time. Firstly, in order to get the approximate existing time of each ancestors on the species tree, we downloaded the dated species tree (Figure S7 in additional file 1) for the 69 species from TIMETREE (www.timetree.org)[71] and use its time information to date our species tree. We dated the ancestral nodes with consistent sub-trees between these two trees. In this way, we got approximate existing time for ancestral nodes where 20 or more families originated. Finally, we dated 23 such ancestral nodes and got the origin time of 9,581 (total: 9,767) gene families.

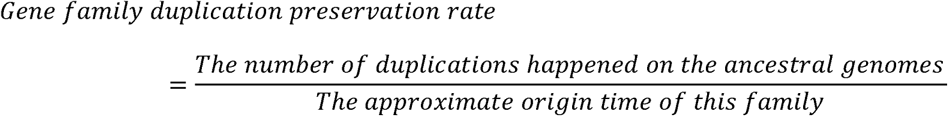

### 4.6.3 Duplication overlap between two branches on species tree

For each ancestral branch, we got a gene family set consisting of gene families expanded at this branch, and we labeled this set as Dl. In this work, we defined a measure named ‘duplication overlap’ to describe the overlap rate of expanded gene families between two branches on the species tree.

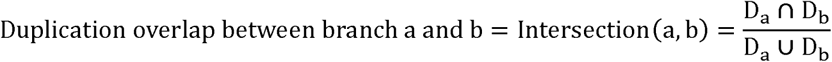

## List of abbreviations

WGD: Whole Genome Duplication
TS: Teleost-Specific
DL: Duplication-Loss
PPI: Protein-Protein Interaction
GO: Gene Ontology
CDS: Sequence coding for amino acids in protein
MSA: multiple sequence alignment
CV: Coefficient of Variance
ML: maximum likelihood

## Acknowledgements

We thank HaiLing Fang and Bing Liu for their assistance in data preparation and figure modification.

## Declarations

### Ethics approval and consent to participate

Not applicable.

### Consent for publication

Not applicable.

### Authors’ contributions

KL conceived of this project and improved the manuscript. JS designed the experiment, performed the analysis and wrote the manuscript. XH downloaded the data and performed some analysis. All authors read and approved the final manuscript.

### Competing interests

The authors have declared no competing interests.

### Funding

This work was supported by the State Key Basic Research and Development Plan (2017YFA0605104) and a project of the State Key Laboratory of Earth Surface Processes and Resource Ecology.

### Availability of data and materials

Data in the study and parameter-learning related scripts are freely available via the website http://cmb.bnu.edu.cn/69vertebrates/.

## Supplementary materials

additional file 1: supplemental figures and tables.

additional file 2: supplemental methods and materials.

