## Supplemental figures and tables for "Evolutionary patterns of 64 vertebrate genomes (species) revealed by phylogenomics analysis of protein-coding gene families"

### Supplementary material

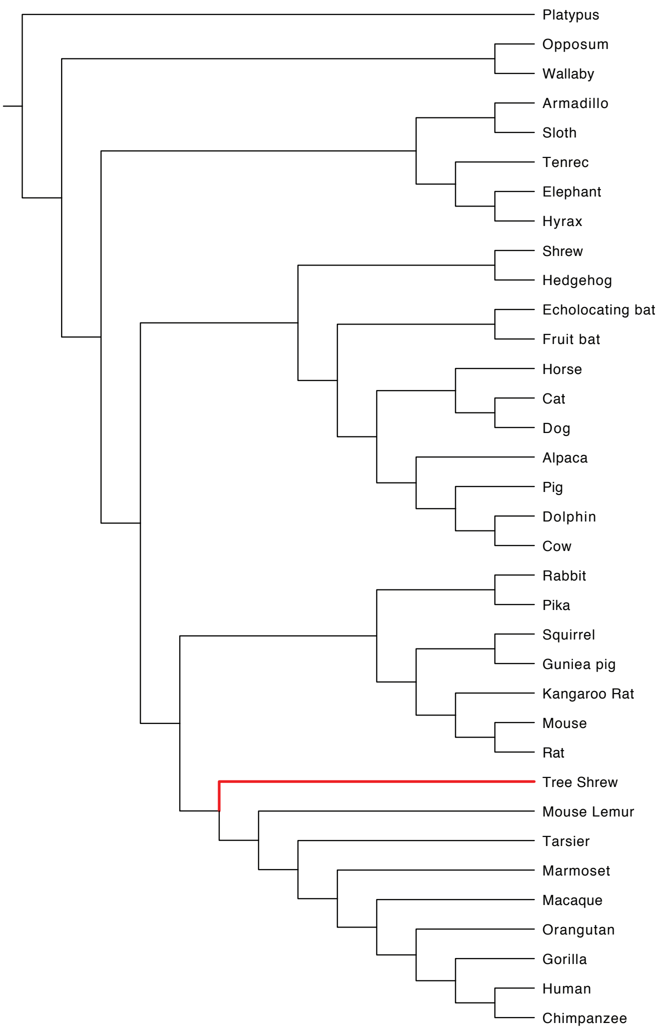


**Figure S1 Mammals' species tree from Song et al. 2012**[1]

The unique incongruence clade between this tree and our final species tree is the Tree Shrew, which labeled in red.

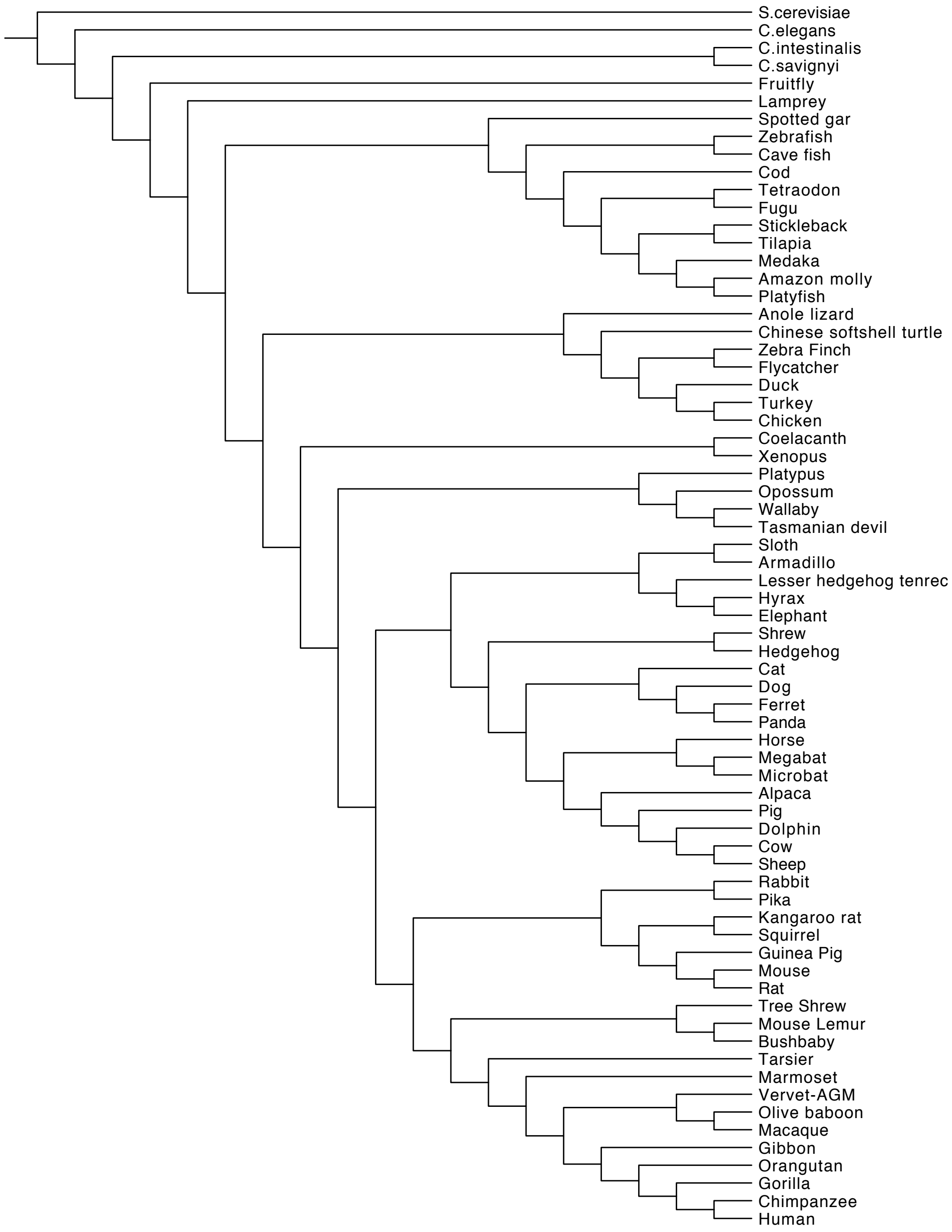


**Figure S2 Species tree inferred by Phyldog**

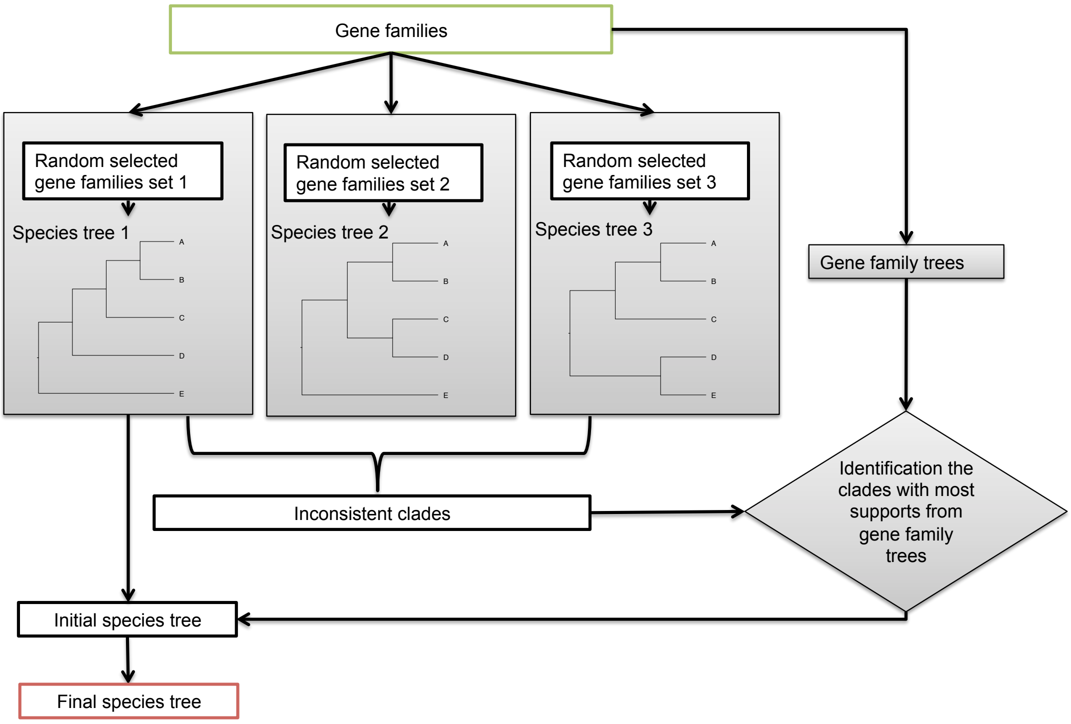


**Figure S3 Alternative species tree inference workflow**

Here, we only randomly selected three gene families sets to infer the initial species tree and we can select more than three sets in real application.

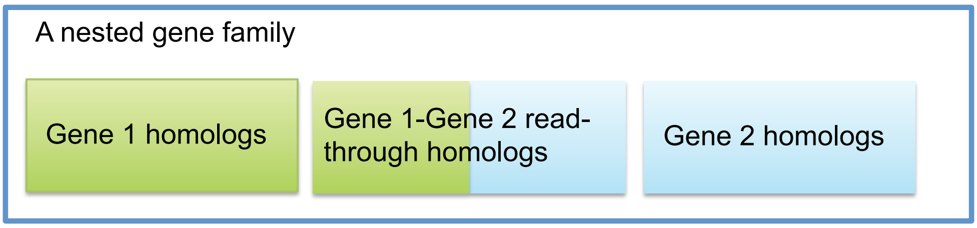


**Figure S4 Diagram of nested gene family**

This diagram displayed the nested gene family caused by read-through genes which read-through two independent genes. And there are more complex nested families, which nested more sub gene families.

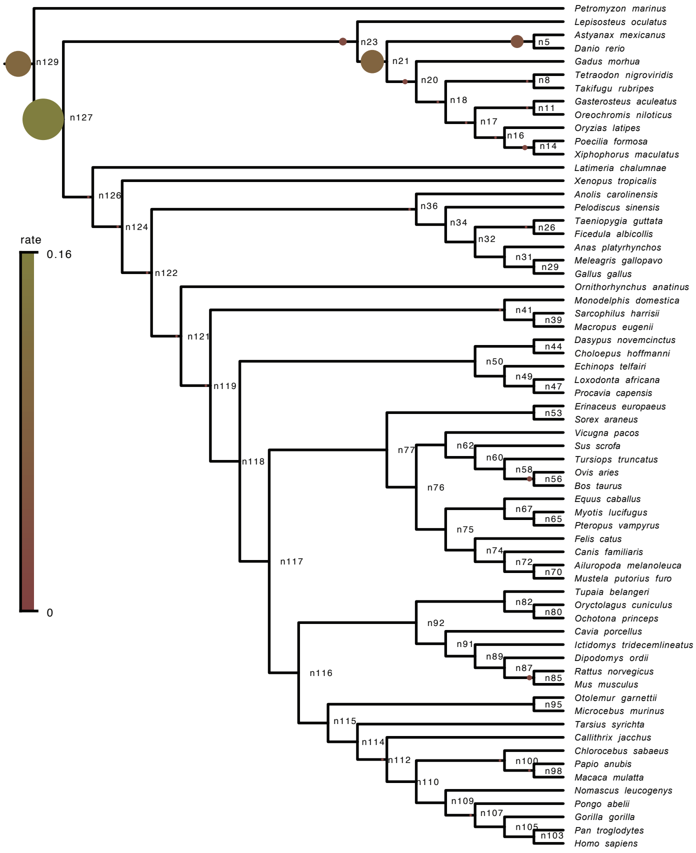


**Figure S5 Average gene duplication retention rates and labeled ancestral nodes**

The size of cycle on each ancestral branch was determined by the average gene duplication retention rates (detailed in Table S1). There are three ancestral branches showing significantly higher average gene duplication rate among these branches (ks.test in R: p-value = 0.005545). These ancestral branches were reported to undergo genome duplication in previous studies[2-6]. This result verified the correctness of our reconciliation results in some ways.

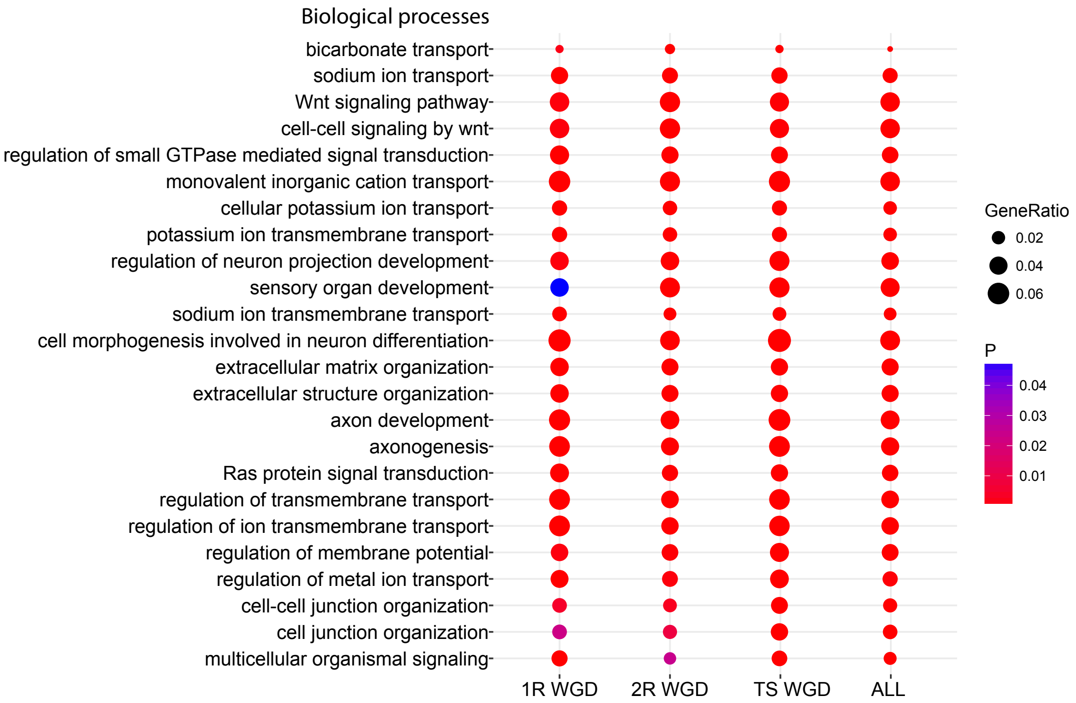


**Figure S6 GO enrichment results of WGD-affected gene families**

Class '1R WGD' represents genes from the gene families with ohnologs retention after the first round WGD in vertebrates. Class '2R WGD' represents genes from the gene families with ohnologs retention after the second round WGD. Class 'TS WGD' represents genes from the gene families with ohnologs retention after the TS WGD. Class 'All' represents genes from gene families with ohnologs retention at least after one of these tree WGDs.

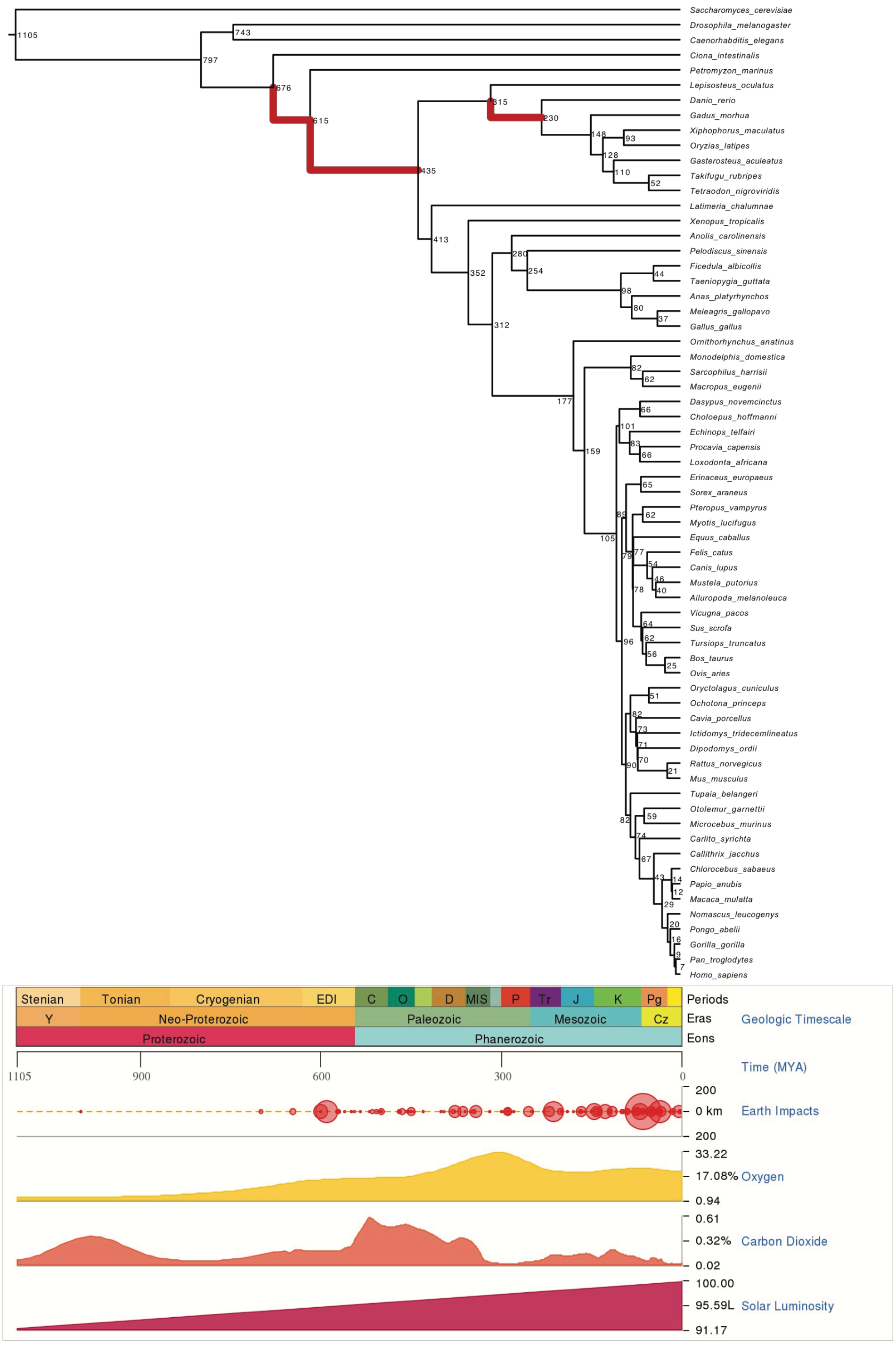


**Figure S7 Dated species tree from TIMETREE**

The concentration of oxygen is displayed under the species tree. The number labeled beside each node represents its age (unit: million years ago, MYA). And the WGD-affected branches are labels in bold and colored in red.

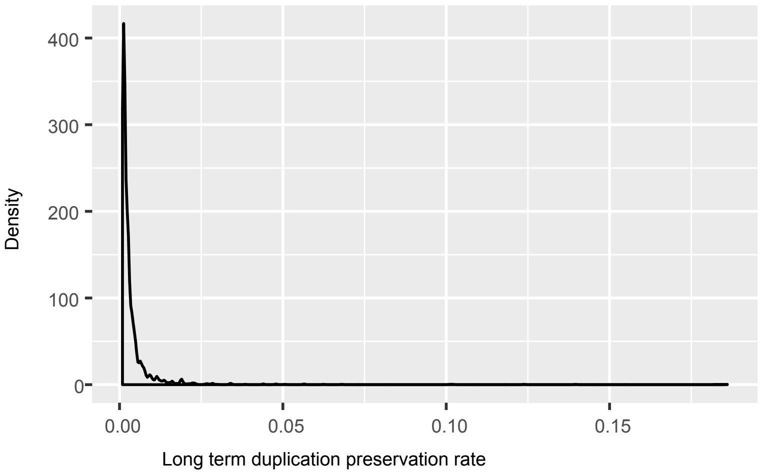


**Figure S8 Distribution of gene family duplication retention/preservation rates with X-axis limited to 0.0009 ~ 0.016**

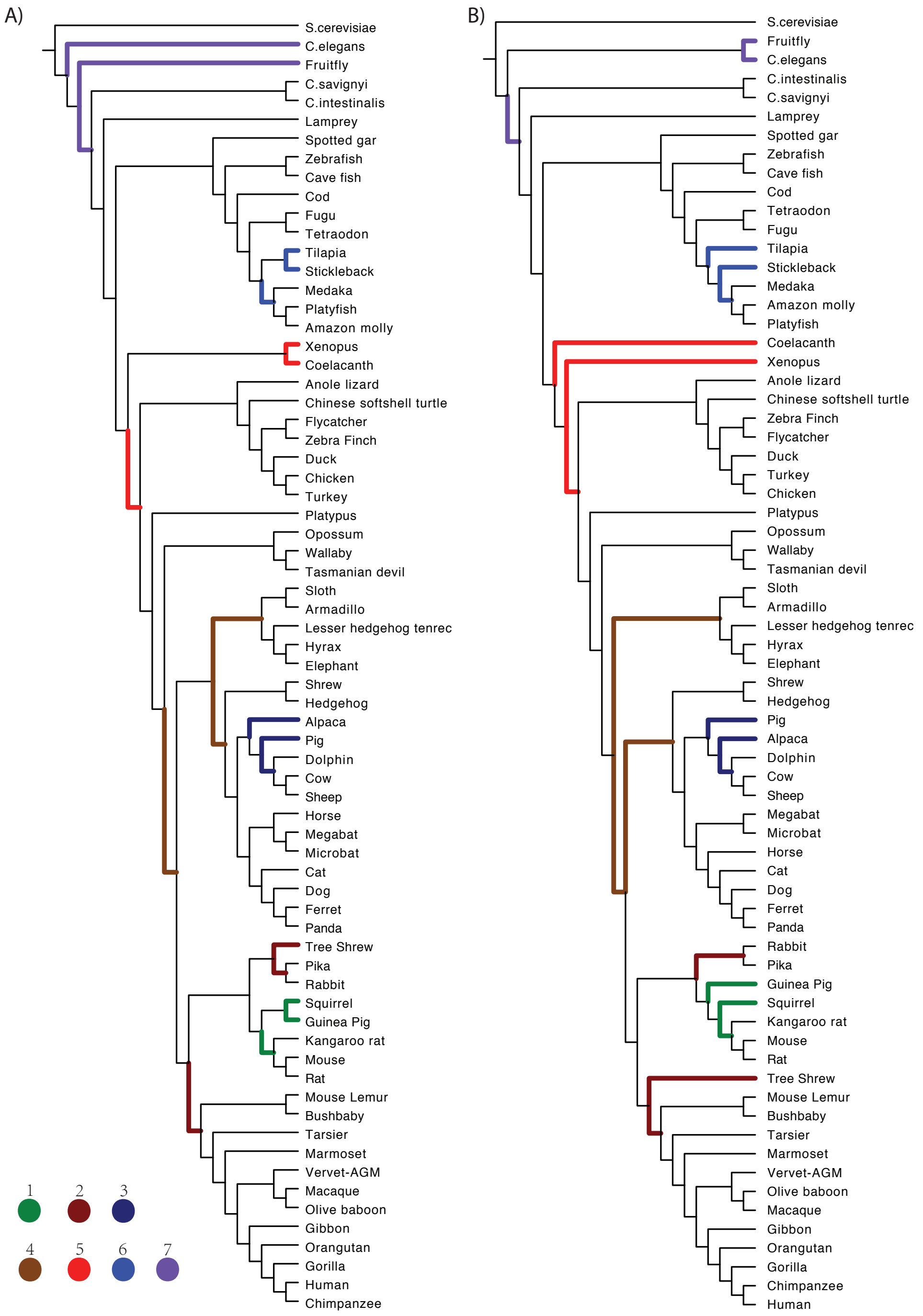


**Figure S9 Inconsistent clades between our initial species tree and Ensembl species tree**

The clades bearing inconsistent phylogenetic relationships between the initial species tree and the Ensembl species tree are labeled in different colors and the inconsistences are numbered. A) Our initial species tree. B) Ensembl species tree.

**Table S1 Average gene duplication retention rate for each branch**

| Branch | Duplication rate | Branch | Duplication rate |
|---|---|---|---|
| *Ornithorhynchus anatinus* | 0.286 | n127 | 0.1573 |
| *Sus scrofa* | 0.2155 | n129 | 0.0998 |
| *Taeniopygia guttata* | 0.1727 | n21 | 0.0863 |
| *Dasypus novemcinctus* | 0.1326 | n5 | 0.0497 |
| *Latimeria chalumnae* | 0.1299 | n23 | 0.0272 |
| *Myotis lucifugus* | 0.1067 | n85 | 0.0217 |
| *Astyanax mexicanus* | 0.1028 | n20 | 0.0209 |
| *Poecilia formosa* | 0.1016 | n14 | 0.0176 |
| *Monodelphis domestica* | 0.0973 | n56 | 0.0175 |
| *Homo sapiens* | 0.0968 | n126 | 0.0133 |
| *Petromyzon marinus* | 0.0946 | n8 | 0.0123 |
| *Callithrix jacchus* | 0.0869 | n41 | 0.0118 |
| *Danio rerio* | 0.0852 | n18 | 0.0111 |
| *Tetraodon nigroviridis* | 0.0829 | n100 | 0.0087 |
| *Anolis carolinensis* | 0.0725 | n121 | 0.0075 |
| *Oryctolagus cuniculus* | 0.0717 | n26 | 0.0075 |
| *Macaca mulatta* | 0.07 | n98 | 0.0075 |
| *Ovis aries* | 0.0643 | n119 | 0.0074 |
| *Otolemur garnettii* | 0.0605 | n11 | 0.0072 |
| *Rattus norvegicus* | 0.0597 | n107 | 0.007 |
| *Xenopus tropicalis* | 0.058 | n124 | 0.0067 |
| *Sarcophilus harrisii* | 0.057 | n17 | 0.006 |
| *Felis catus* | 0.056 | n112 | 0.0058 |
| *Pelodiscus sinensis* | 0.0523 | n122 | 0.0058 |
| *Lepisosteus oculatus* | 0.0496 | n16 | 0.0054 |
| *Ictidomys tridecemlineatus* | 0.0485 | n36 | 0.0052 |
| *Oreochromis niloticus* | 0.048 | n47 | 0.0048 |
| *Gorilla gorilla* | 0.0479 | n103 | 0.0047 |
| *Echinops telfairi* | 0.0473 | n29 | 0.0046 |
| *Cavia porcellus* | 0.0471 | n72 | 0.0046 |

| Branch | Duplication rate | Branch | Duplication rate |
|---|---|---|---|
| *Anas platyrhynchos* | 0.047 | n118 | 0.0045 |
| *Mus musculus* | 0.0456 | n44 | 0.0045 |
| *Oryzias latipes* | 0.044 | n91 | 0.0045 |
| *Loxodonta africana* | 0.0437 | n31 | 0.0044 |
| *Canis familiaris* | 0.0416 | n75 | 0.0041 |
| *Gadus morhua* | 0.0391 | n110 | 0.004 |
| *Gasterosteus aculeatus* | 0.0386 | n34 | 0.004 |
| *Equus caballus* | 0.0376 | n74 | 0.0038 |
| *Meleagris gallopavo* | 0.0364 | n60 | 0.0036 |
| *Takifugu rubripes* | 0.0343 | n32 | 0.0034 |
| *Mustela putorius furo* | 0.0317 | n105 | 0.0033 |
| *Pongo abelii* | 0.0301 | n70 | 0.0032 |
| *Ailuropoda melanoleuca* | 0.0267 | n109 | 0.0031 |
| *Gallus gallus* | 0.0262 | n53 | 0.0031 |
| *Tupaia belangeri* | 0.0247 | n58 | 0.0031 |
| *Nomascus leucogenys* | 0.0229 | n80 | 0.003 |
| *Xiphophorus maculatus* | 0.0225 | n76 | 0.0025 |
| *Erinaceus europaeus* | 0.0222 | n95 | 0.0025 |
| *Bos taurus* | 0.0214 | n89 | 0.0022 |
| *Ochotona princeps* | 0.0201 | n49 | 0.002 |
| *Ficedula albicollis* | 0.02 | n117 | 0.0018 |
| *Sorex araneus* | 0.0179 | n67 | 0.0014 |
| *Choloepus hoffmanni* | 0.0173 | n39 | 0.0013 |
| *Tarsius syrichta* | 0.0155 | n115 | 0.0012 |
| *Microcebus murinus* | 0.0152 | n65 | 0.001 |
| *Macropus eugenii* | 0.0151 | n116 | 0.0009 |
| *Chlorocebus sabaeus* | 0.015 | n87 | 0.0008 |
| *Papio anubis* | 0.0139 | n114 | 0.0005 |
| *Dipodomys ordii* | 0.0114 | n50 | 0.0005 |
| *Tursiops truncatus* | 0.0098 | n77 | 0.0005 |
| *Procavia capensis* | 0.0091 | n62 | 0.0003 |

| Branch | Duplication rate | Branch | Duplication rate |
|---|---|---|---|
| *Pan troglodytes* | 0.0089 | n82 | 0.0003 |
| *Pteropus vampyrus* | 0.0066 | n92 | 0.0003 |
| *Vicugna pacos* | 0.0064 | | |

In this table, ancestral branches are represented by their end-nodes. And the ancestral node represents the ancestral vertebrate species as the Figure S5 displayed. The calculation methods are detailed in section 3 in this file.

2R & TS WGD: Gene families retaining ohnologs both from the TS and 2R WGDs.

1R & 2R & 3R: Gene families retaining ohnologs from all of the three WGDs.

#### Table S2 PPI enrichment results in the WGD-affected gene family classes

| WGDs | PPI | Gene pairs | Enrichment fold | P-value |
|---|---|---|---|---|
| 1R WGD | 8,340 | 1,214,080 | 4.245023 | < 2.2e-16 |
| 2R WGD | 15,690 | 4,179,953 | 2.369621 | < 2.2e-16 |
| TS WGD | 11,838 | 2,182,655 | 3.407453 | < 2.2e-16 |
| 1R & 2R WGD | 1,096 | 340,918 | 1.880676 | < 2.2e-16 |
| 1R & TS WGD | 2,746 | 446,232 | 3.656845 | < 2.2e-16 |
| 2R & TS WGD | 3,144 | 1,190,374 | 1.552143 | < 2.2e-16 |
| 1R & 2R & 3R | 428 | 76,538 | 3.273032 | < 2.2e-16 |

1R WGD: Gene families that only retaining ohnologs after the first round WGD.

2R WGD: Gene families that only retaining ohnologs after the second round WGD.

TS WGD: Gene families that only retaining ohnologs after the TS (teleost fishes) specific WGD.

1R & 2R WGD: Gene families retaining ohnologs both from the 1R and 2R WGDs.

1R & TS WGD: Gene families retaining ohnologs both from the TS and 1R WGDs

#### Table S3 Supports of the 7 inconsistent clades from 11,698 gene family trees

| Inconsistent clades[a] | Ensembl species tree | Guenomu species tree |
|---|---|---|
| Inconsistent clade 1 | 2066* | 1332 |
| Inconsistent clade 2 | 1592 | 2172* |
| Inconsistent clade 3 | 672 | 2932* |
| Inconsistent clade 4 | 2650* | 147 |
| Inconsistent clade 5 | 2834* | 676 |

| Inconsistent clades[a] | Ensembl species tree | Guenomu species tree |
|---|---|---|
| Inconsistent clade 6 | 1731 | 2752* |
| Inconsistent clade 7 | 501 | 405 |

*The favored clade

[a] The relative clades can be found in Figure S9
