## Supplemental notes for "Evolutionary patterns of 64 vertebrate genomes (species) revealed by phylogenomics analysis of protein-coding gene families"

### Gene family trees inference

After the gene family classification, we got totally 54,808 gene families. There were 17,888 gene families including more than one member and 17,572 gene families including members from more than one species. Then, the gene families with members from a unique species were filtered out from our data. Further, gene families including known read-through genes annotated on human gene annotation file (v24) in ENCODE[1] were removed. The gene families with very members more than 1,000 were also removed and 17,025 gene families were left for following analysis. In this step, protein sequences of 17,025 gene families were aligned in MAFFT v7[2](--auto) and then translated into CDS alignments by translatorX[3]. The poorly aligned regions were removed from these CDS MSAs by trimAl[4]. Then, we removed some gene families with specific labels in its sequences (such as X) or with very poor alignment quantity. More details, there were 63 gene families MSA containing specific labels, two gene families got too poor MSA quantity that trimAl refused to trim and 2,923 families received MSAs less than 100bp after trim. For the left 14,037 CDS MSAs, we inferred the gene family tree in RAxML v8.2.9[5] under GTRGAMMI sequence evolution model. For some MSAs including members less than four and some MSAs including too much gaps, so we finally got reliable phylogenetic trees for 11,698 gene families only. Finally, we inferred optimal parameters set and rearranged gene family tree for each gene families. There were 1,917 gene families couldn't obtain consistent optimal parameters pairs according to the parameter-learning process and 14 gene families got consistent optimal parameters pairs but inconsistent reconciled results under these parameters pairs.

Above all, we finally got 9,767 duplication/loss annotated gene family trees as our core results.

### Reconciliation

In this study, we mainly used 15 pairs of (costdup, costloss) with different 'costdup/costloss' ratios to finish the reconciliation and parameter-learning process. And the pairs are:

(1,1), (2,1), (1,2), (2,3), (3,2), (1,3), (3,1), (1,4), (4,1), (1,8), (8,1), (1,10), (10,1), (1,12), (12,1)

### Average gene duplication retention rate for each branch

Based on our reconciliation results (9,767 gene families), we calculated the average gene duplication retention rate for each branch in the vertebrates' part (63 branches) between every two successive ancestral species nodes on the species tree.

$$Average\ gene\ duplication\ retention\ rate\ for\ each\ branch = \frac{\sum relative\ gene\ duplication\ rate\ on\ this\ branch\ of\ each\ gene\ family}{the\ number\ of\ gene\ families\ that\ containing\ members\ on\ this\ branch}$$

Among them

$$The\ relative\ gene\ duplication\ of\ a\ specific\ gene\ family\ on\ a\ specific\ branch = \frac{the\ number\ of\ duplication\ events\ of\ this\ gene\ family\ on\ this\ branch}{the\ gene\ family\ size\ of\ this\ gene\ family\ at\ the\ beginning\ of\ this\ branch}$$

1313.
